# The Anti-Inflammatory Role of GPNMB in Post-Traumatic Osteoarthritis

**DOI:** 10.1101/2025.06.06.658389

**Authors:** Asaad A. Al-Adlaan, Bryson Cook, Nazar J. Hussein, Fatima A. Jaber, Trinity A. Kronk, Ernesto Solorzano Z., Salvatore Frangiamore, Hope C. Ball, Fayez F. Safadi

## Abstract

Osteoactivin (GPNMB) is a transmembrane protein expressed in multiple cell types with known functions in muscle, bone, and neurons, but the role of GPNMB in chondrocytes and cartilage homeostasis remains unknown. Here we show *GPNMB/Gpnmb* is expressed in human and mouse primary chondrocytes, and that its expression is increased in damaged human cartilage and under pro-inflammatory conditions. We report that recombinant GPNMB treatment inhibits the expression of *Mmps* (*Mmp3, 9*, and *13*), *Adamts4* and *Il6* following IL-1β-stimulation *in vitro. In vivo*, GPNMB function was assessed in a post-traumatic osteoarthritis model, destabilization of the medial meniscus (DMM). Transgenic animals lacking functional GPNMB protein (DBA/2J) developed severe cartilage damage and demonstrated significant increases in pro-inflammatory cytokine expression following DMM. To elucidate the mechanism of action, we demonstrate that GPNMB regulates the MAPK signaling pathway via pERK inhibition in primary murine chondrocytes. Taken together, our results identify a novel anti-inflammatory role for GPNMB in cartilage and chondrocytes and identify GPNMB as a potential therapeutic modality for inflammatory joint diseases.

## Introduction

Osteoactivin, also known as glycoprotein non-metastatic melanoma protein B (GPNMB), is a transmembrane glycoprotein first identified in a rat model of osteopetrosis (Safadi et al., 2002). GPNMB has subsequently been found in many cell types including osteoblasts, osteoclasts, macrophages, hepatocytes, dendritic cells and cancer cells and is known to be involved in a plethora of functions including proliferation, differentiation, migration, angiogenesis, osteogenesis, and apoptosis (Arosarena et al., 2018; Rose et al., 2010; Schwarzbich et al., 2012; Selim et al., 2003; Sheng et al., 2008; Yu et al., 2016). GPNMB encodes a 574 amino acid protein that includes an arginine-glycine-aspartic acid (RGD) domain, a polycystic kidney disease domain (PKD) and a proline rich repeat (PRRD) domain (**Figure 1-Figure Supplement 1**) (Owen et al., 2003). The majority of the biological function, however, is conducted by the extracellular domain of the protein, shed from the surface by furin-like proteases and extracellular proteases, such as a disintegrin and metalloproteinase with thrombospondin motifs-10 (Adamts-10) (Kawahara et al., 2016; Murata et al., 2015; Oyewumi et al., 2016; Rose et al., 2010).

The role of GPNMB in fracture repair and bone regeneration is well studied. Upregulated in response to skeletal damage, GPNMB promotes bone formation through either direct or indirect regulation of osteoblast proliferation and adhesion through multiple signaling pathways (Abdelmagid et al., 2010; Abdelmagid et al., 2008; S. M. Abdelmagid et al., 2015; Frara et al., 2016). GPNMB also functions as a negative regulator of osteoclast differentiation and survival (S. M. Abdelmagid et al., 2015; Sheng et al., 2008; Sondag et al., 2016). Indeed, transgenic GPNMB mice exhibit increased bone volume, mineralization and thickness, increased osteoblast number and decreased osteoclast survival compared to C57BL/6 control animals (S. M. Abdelmagid et al., 2015; Frara et al., 2016; Sondag et al., 2016).

GPNMB is emerging as a crucial factor in the inflammatory response following tissue injury (Li et al., 2010). GPNMB levels were elevated in both mouse and rat models after myocardial infarction (Järve et al., 2017). GPNMB has also been shown to have an anti-inflammatory role in macrophages, reducing the expression of pro-inflammatory signaling molecules such as interleukin-6 (IL-6), interleukin-12p40 (IL-12p40) and nitric oxide (NO) (Li et al., 2010; Ripoll et al., 2007; Zhou et al., 2017). Furthermore, GPNMB has been shown to play a role in the synthesis of extracellular matrix (ECM) components and its expression is increased with inflammatory stimuli and ECM damage, conditions common in the pathophysiology of degenerative joint diseases such as osteoarthritis (OA) (Jiang et al., 2011; Owen et al., 2003; Safadi et al., 2002; Sun et al., 2015). Despite the above roles, the biological function of GPNMB in normal or damaged cartilage and chondrocyte homeostasis have not yet been studied.

To address this knowledge gap, we first determined *GPNMB* expression in human primary chondrocytes and cartilage. *GPNMB* expression was increased in damaged cartilage from osteoarthritic patients compared to normal (no history of rheumatic disease) and following pro-inflammatory stimulus in human primary chondrocytes and cell-lines. Furthermore, recombinant GPNMB treatment significantly lowered expression of catabolic markers in human primary chondrocytes and reduced ECM degradation in cartilage explants. We subsequently validated these results in murine primary chondrocytes and examined endogenous *Gpnmb* expression in a mouse model: DBA/2J mice, which produce a truncated and non-functional GPNMB protein (Abdelmagid et al., 2014). To better understand the role of GPNMB *in vivo*, we induced post-traumatic OA via destabilization of the medial meniscus (DMM) surgery in DBA/2J mice (Culley et al., 2015; Glasson et al., 2007). The results demonstrated functional GPNMB is essential for the reduction of inflammation and protection against post-traumatic cartilage damage. Collectively, our results are the first to show the extracellular domain of GPNMB serves an anti-inflammatory function in primary chondrocytes and cartilage and reduces catabolic gene expression both *in vitro* and *in vivo* and suggests GPNMB may be a potential anti-inflammatory therapy for inflammatory joint diseases.

## Results

### *GPNMB* expression is elevated in damaged human cartilage

We first assessed the expression levels of *GPNMB* in human OA cartilage samples. To confirm we had accurately harvested cartilage from regions of both low- and high-grade OA damage, we began by first macroscopically evaluating the cartilage using India Ink, where particles adhere to fibrillated tissue indicative of cartilage damage (Schmitz et al., 2010). Cartilage was isolated from low- and high-grade cartilage samples for mRNA analyses. We examined expression levels of a key cartilage ECM component, aggrecan, encoded by the *ACAN* gene. Expression levels of *ACAN* were significantly decreased in high-grade OA cartilage compared to low-grade regions supporting our assessment of cartilage damage (**Figure 1A**). We then determined expression levels of *GPNMB* in cartilage harvested from the low-grade and high-grade OA cartilage. Results demonstrated a significant increase in *GPNMB* expression in high-grade cartilage versus low-grade cartilage levels (**Figure 1B**). This data and previous findings regarding the anti-inflammatory role of GPNMB in other cell types led us to postulate that GPNMB has an anti-inflammatory role in primary chondrocytes.

**Figure 1:**
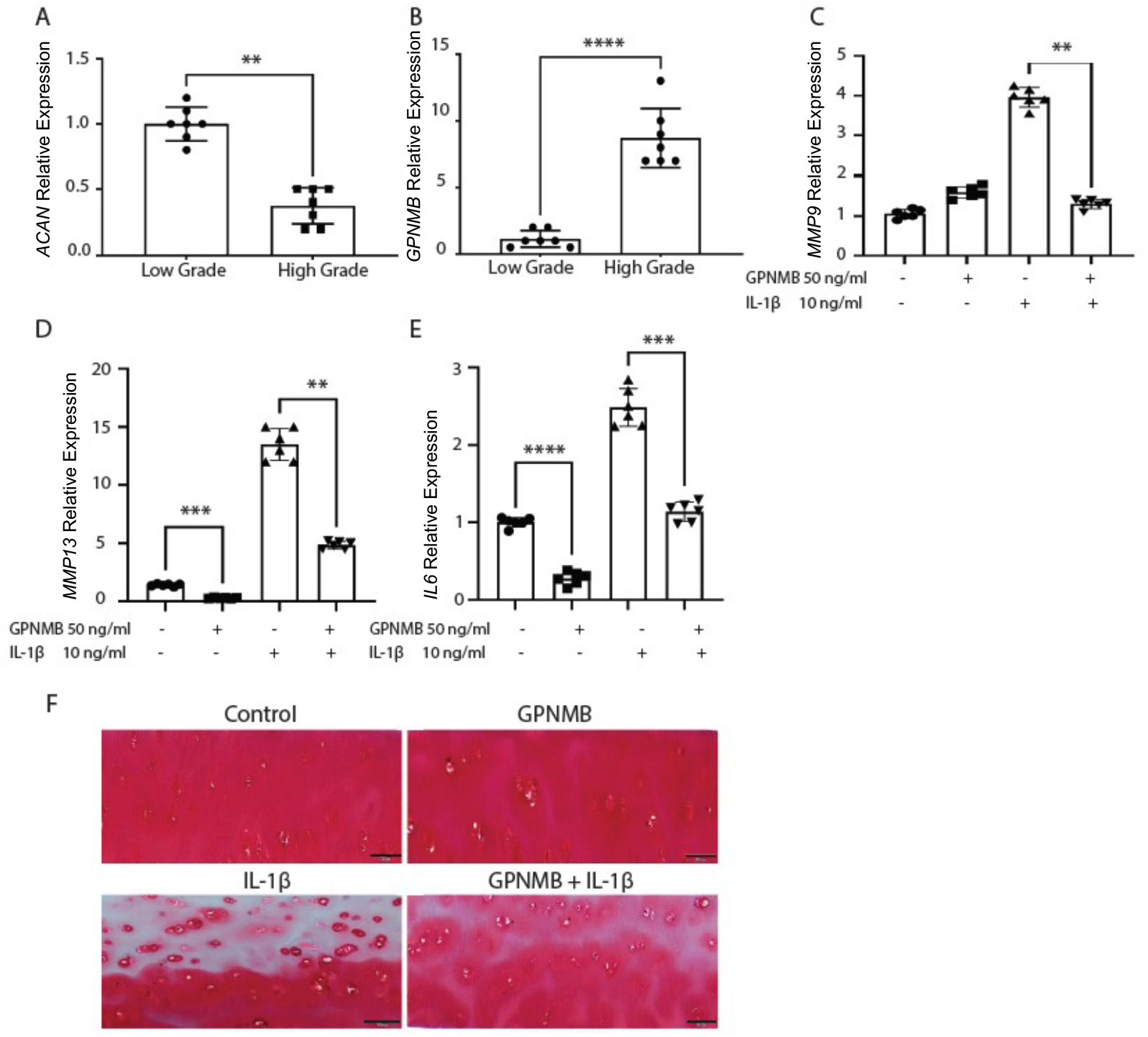
*GPNMB* expression is increased in high-grade OA cartilage and recombinant GPNMB reduces catabolic gene expression in human HTB-94 cell culture and cartilage damage in human cartilage explants. Human knee articular cartilage was isolated from low-grade and high-grade OA joints and mRNA expression of (**A**) *ACAN* and (**B**) *GPNMB* were determined via RT-qPCR. HTB-94 cells treated with combination of recombinant GPNMB and IL-1β demonstrated significantly decreased expression of (**C**) *MMP9* (**D**) *MMP13* and (**E**) *IL6* mRNAs. Combined recombinant GPNMB and IL-1β treatment significantly reduced cartilage degradation in human explant cultures compared to IL-1β treatment alone in Safranin-O-stained human cartilage explants (**F**). Scale bar= 200µm in top panels and 100µm in magnified lower images. Data presented as Mean ± SEM (**A-E**: n=7, 3 replicates per experiment; **F**: Experiments were repeated three times, 3 replicates per experiment with similar results. Representative images are shown). **= p<0.01, ***=p<0.001, ****=p<0.0001.

### GPNMB treatment reduces catabolic gene expression in IL-1β stimulated HTB-94 chondrocytes

One of the hallmarks of OA is increased inflammation within the joint and an increase in the production of inflammatory chemokines and cytokines such as interleukin-1 beta (IL-1β), interleukin-6 (IL-6), matrix metalloproteinase-9 (MMP-9) and matrix metalloproteinase -13 (MMP-13) (Gauci et al., 2017; Goldring, 2012; Goldring & Goldring, 2004; Robinson et al., 2016). Since *GPNMB* was shown to be elevated in more damaged regions of human cartilage, we theorized that increased GPNMB levels may serve a protective anti-inflammatory role. To test this, we utilized human chondrosarcoma cells (HTB-94) which maintain a chondrocyte phenotype in culture and have been established as an appropriate model for examinations of inflammation and inflammatory signaling pathways (Mengshol et al., 2001). Cells were serum starved overnight and then either left untreated or treated for 24 hours with either recombinant GPNMB alone, IL-1β alone or a combination of recombinant GPNMB and IL-1β. As expected, IL-1β treatment significantly increased expression of *MMP9, MMP13* and *IL6* (**Figure 1C-E**). Interestingly, treatment with recombinant GPNMB alone significantly decreased gene expression of both *MMP13* and *IL6* and inclusion of GPNMB with IL-1β reduced catabolic expression of all target genes (**Figure 1C-E**). Together, this suggests GPNMB has an anti-inflammatory role in human primary chondrocytes.

### Human explant cultures treated with GPNMB display low ECM degradation after IL-1β stimulation

Having determined catabolic gene expression was reduced with recombinant GPNMB treatment in HTB-94 chondrocytes, we next determined if GPNMB pharmacological treatment could reduce ECM degradation following IL-1β stimulation *ex vivo* in human explant cultures. Undamaged human explants were serum starved overnight then left untreated or treated for 24 hours with recombinant GPNMB alone, IL-1β alone or a combination of recombinant GPNMB and IL-1β. Safranin-O-stained explant results showed severe ECM degradation in cultures treated with IL-1β alone compared to untreated controls (**Figure 1F**). Additionally, the severity of the degradation was reduced with the inclusion of recombinant GPNMB (**Figure 1F**). These results establish that recombinant GPNMB treatment decreases IL-1β-induced inflammation in human chondrocytes and cartilage.

### IL-1β stimulation induces *Gpnmb* expression in murine primary chondrocytes

Examinations of endogenous expression of *Gpnmb* in human cartilage and chondrocytes demonstrated that *Gpnmb* expression was upregulated in high-grade damaged OA cartilage and following IL-1β proinflammatory stimulation in human primary chondrocytes. Next, we confirmed these results in murine primary chondrocytes. Primary chondrocytes were isolated from 5-6 day old pups (n=6) as previously described and serum starved overnight (Gosset et al., 2008; Jonason et al., 2015). The chondrocytes were then treated, in triplicate, with either regular culture media (control) or IL-1β for 24 hours and the expression of *Gpnmb* was determined via RT-qPCR. Results showed a significant increase in *Gpnmb* expression following IL-1β stimulation compared to untreated controls (**Figure 2A**).

**Figure 2:**
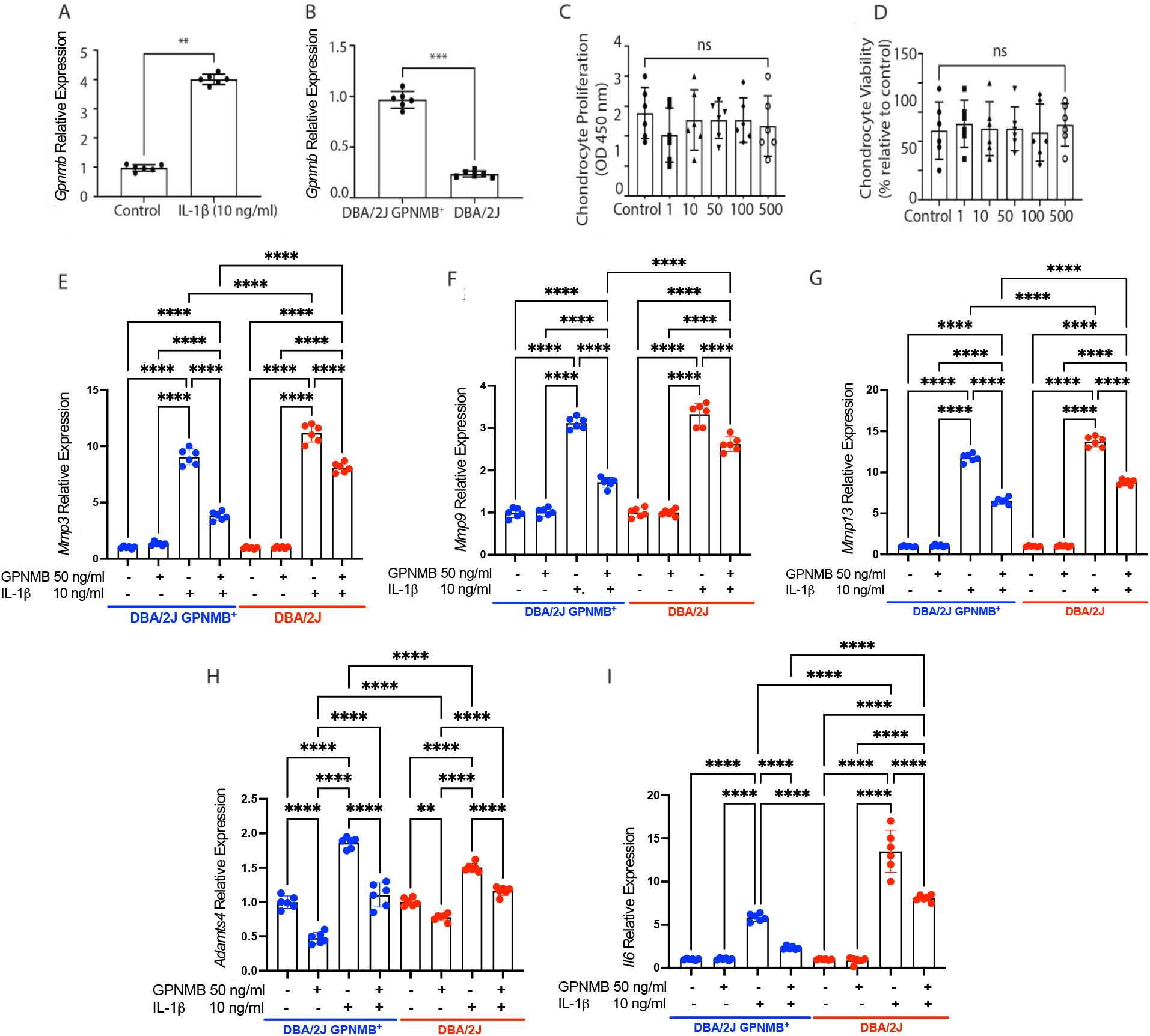
Gpnmb expression is induced with IL-1β proinflammatory stimulation, reduced in DBA/2J mice and recombinant GPNMB inhibits catabolic expression in DBA/2J GPNMB^+^ and DBA/2J chondrocytes. (**A**) *Gpnmb* expression is increased with pro-inflammatory IL-1β treatment. *Gpnmb* expression (**B**) is significantly reduced in DBA/2J mice. Recombinant GPNMB treatment does not affect chondrocyte proliferation (**C**) or viability (**D**). Recombinant GPNMB inhibits *Mmp3* (**E**), *Mmp9* (**F**), *Mmp13* (**G**) expression in DBA/2J GPNMB^+^ and DBA/2J chondrocytes and reduces *Adamts4* (**H**) and *Il6* (**I**) expression. Groups are compared amongst their own genotype and the associated treatment group of the other genotype with only significant results shown (**E-I)**. Scale bar= 200µm. Data presented as Mean ± SEM (n=6-8, 3 replicates per experiment). *=p<0.05, **= p<0.01, ***=p<0.001, ****=p<0.0001.

### Endogenous *Gpnmb* is reduced in DBA/2J mutant mice

To study the role of GPNMB in chondrocytes, we utilized a mouse model with a natural mutation in GPNMB (DBA/2J). This global mouse model produces a truncated and non-functional GPNMB protein (Abdelmagid et al., 2014). For this experiment, we determined endogenous *Gpnmb* levels in primary chondrocytes of DBA/2J GPNMB^+^ (control) and DBA/2J mice. Results revealed *Gpnmb* mRNA levels in the mutant DBA/2J mice (n=6) were significantly lower than those found in DBA/2J GPNMB^+^ (n=6) chondrocytes (**Figure 2B**). Our results are consistent with the literature which shows that that the *Gpnmb* gene in DBA/2J mice carries a nonsense mutation that leads to reduced RNA stability (Anderson et al., 2008).

### Recombinant GPNMB treatment does not affect murine chondrocyte viability or proliferation

Before investigating the effects of GPNMB on murine chondrocytes *in vitro*, we first determined if pharmacological treatment with recombinant GPNMB would affect primary chondrocyte viability or proliferation. Chondrocytes were cultured with increasing doses of GPNMB for 72 hours, and cell viability and proliferation were assessed with MTT and CyQuant assays, respectively. Data revealed no significant changes in chondrocyte viability or proliferation with increasing concentrations of recombinant GPNMB (**Figure 2C, D**). These results suggest that treatment has no adverse or toxic effects on chondrocytes.

### GPNMB treatment inhibits *Mmp, Adamts4* and *Il6* expression in DBA/2J GPNMB^+^ and DBA/2J chondrocytes

To determine if treatment with recombinant GPNMB would alter catabolic gene expression in murine primary chondrocytes, we isolated femoral and tibial articular cartilage from 5-6 day old male control DBA/2J GPNMB^+^ (n=6) and mutant DBA/2J mouse pups (n=6) and plated the primary chondrocytes of each mouse model in triplicate. Before treatment, we examined endogenous mRNA levels of *Mmp3, Mmp9* and, *Mmp13* in control DBA/2J GPNMB^+^ and DBA/2J primary chondrocytes. Results found no significant differences in endogenous mRNA expression in these strains (**Figure 2-Figure Supplement 2**). The isolated chondrocytes were serum starved overnight and either left untreated (control) or treated with recombinant GPNMB alone, IL-1β alone or a combination of IL-1β and recombinant GPNMB for 24 hours. Expression levels of cartilage-degrading metalloproteinases *Mmp3, Mmp9* and *Mmp13* were determined via RT-qPCR analysis.

In DBA/2J GPNMB^+^ mice, results showed mRNA levels of *Mmp3, Mmp9* and *Mmp13* were not significantly different between untreated chondrocytes and those treated with recombinant GPNMB alone (**Figure 2E-G**). Whereas chondrocytes treated with IL-1β alone showed significant upregulation of *Mmp3, Mmp9*, and *Mmp13* (**Figure 2E-G**). Interestingly, when DBA/2J GPNMB^+^ chondrocytes were treated with a combination of IL-1β and recombinant GPNMB results demonstrated expression of all three catabolic markers were significantly reduced, suggesting that GPNMB might play a role as an anti-inflammatory factor in chondrocytes (**Figure 2E-G**).

Mutant DBA/2J primary chondrocyte results showed no significant differences in *Mmp3, Mmp9* or *Mmp13* expression between untreated cells and those treated with GPNMB alone (**Figure 2E-G**). Chondrocytes treated with IL-1β alone demonstrated significant increases in *Mmp3, Mmp9* and *Mmp13* mRNA levels (**Figure 2E-G**). Interestingly, these increases in *Mmp3* and *Mmp13* were significantly higher in DBA/2J chondrocytes compared the chondrocytes from DBA/2J GPNMB^+^ controls (**Figure 2E-G**). Primary DBA/2J chondrocytes treated with the combination of IL-1β and recombinant GPNMB showed significant reduction in IL-1β-stimulated increased in *Mmp3, Mmp9* and *Mmp13* expression (**Figure 2E-G**). Together, these results indicate GPNMB has anti-inflammatory functions and reduces expression of IL-1β-stimulated *Mmps*.

In addition to MMPs, which target and degrade ECM collagens, we also evaluated whether recombinant GPNMB would affect expression of *Il6* and an ACAN degrading enzyme, *Adamts4*, in DBA/2J GPNMB^+^ and DBA/2J primary chondrocytes. First, we evaluated endogenous levels of *Adamts4* and *Il6* in DBA/2J GPNMB^+^ and DBA/2J primary chondrocytes and found no significant differences in mRNA expression between the strains (**Figure 2-Figure Supplement 2**). Then, to study the effects recombinant GPNMB treatment on *Adamts4* and *Il6* expression, primary chondrocytes from both strains were serum starved overnight prior to treatment with recombinant GPNMB alone, IL-1β alone or a combination of IL-1β and recombinant GPNMB for 24 hours.

First, we examined *Adamts4* mRNA expression in DBA/2J GPNMB^+^ chondrocytes (n=6). Results showed that treatment with recombinant GPNMB alone was sufficient to significantly reduce *Adamts4* mRNA expression in treated chondrocytes compared to untreated controls (**Figure 2H**). Treatment with IL-1β alone significantly increased expression of *Adamts4*, while treatment with recombinant GPNMB and IL-1β significantly reduced *Adamts4* expression in DBA/2J GPNMB^+^ chondrocytes (**Figure 2H**). In chondrocytes derived from DBA/2J mice (n=6), treatment with recombinant GPNMB alone significantly reduced *Adamts4* expression compared to untreated controls, while IL-1β treatment alone significantly increased *Adamts4* expression (**Figure 2H**). Treatment with the combination of recombinant GPNMB and IL-1β significantly reduced *Adamts4* levels in treated chondrocytes (**Figure 2H**).

Next, we assessed mRNA expression of *Il6* in primary chondrocytes isolated from DBA/2J GPNMB^+^ and DBA/2J mice (n=6, each strain). In both strains, recombinant GPNMB treatment alone resulted in no significant difference in *Il6* mRNA compared to expression untreated controls (**Figure 2I**). Similarly, IL-1β treatment alone significantly increased *Il6* expression in chondrocytes of both mouse strains (**Figure 2I**). Results also revealed that IL-1β-induced expression of proinflammatory *Il6* was significantly higher (a 9-fold) in mutant DBA-2J mice than in DBA/2J GPNMB^+^ control animals. Furthermore, when treated with IL-1β and recombinant GPNMB, *Il6* expression was significantly reduced in DBA/2J GPNMB^+^ chondrocytes (4-fold) and in those from DBA/2J (9-fold) mice with reduction significantly higher than those of controls (**Figure 2I**). Together, these results indicate GPNMB has an anti-inflammatory effect in murine primary chondrocytes by reducing expression of both *Adamts4* and *Il6*.

### Recombinant GPNMB treatment does not alter anabolic gene expression

Having determined that recombinant GPNMB treatment lowered IL-1β-induced catabolic gene expression in human and mouse primary chondrocytes, we next assessed if GPNMB would affect anabolic gene expression. We began by first determining endogenous mRNA levels of anabolic genes SRY-Box Transcription Factor 9 (*Sox9*), type-II collagen (*Col2a1*) and *Acan* in DBA/2J GPNMB^+^ and DBA/2J chondrocytes. Results determined no significant difference in gene expression (**Figure 3-Figure Supplement 3**). To determine the effect of recombinant GPNMB on primary chondrocytes, the cells were serum starved overnight and then treated for 24 hours with either media (control), recombinant GPNMB alone, IL-1β alone or a combination of IL-1β and recombinant GPNMB. mRNA expression was assessed via RT-qPCR.

Results in DBA/2J GPNMB^+^ chondrocytes (n=6) showed recombinant GPNMB treatment alone did not significantly alter anabolic gene expression compared to untreated control cells (**Figure 3A, B, C**). Chondrocytes treated with IL-1β alone demonstrated a significant decrease in *Sox9, Col2a1* and *Acan* mRNA expression **(Figure 3A, B, C**). Interestingly, the combined treatment of recombinant GPNMB and IL-1β showed no significant effects on anabolic gene expression (**Figure 3A, B, C**).

**Figure 3:**
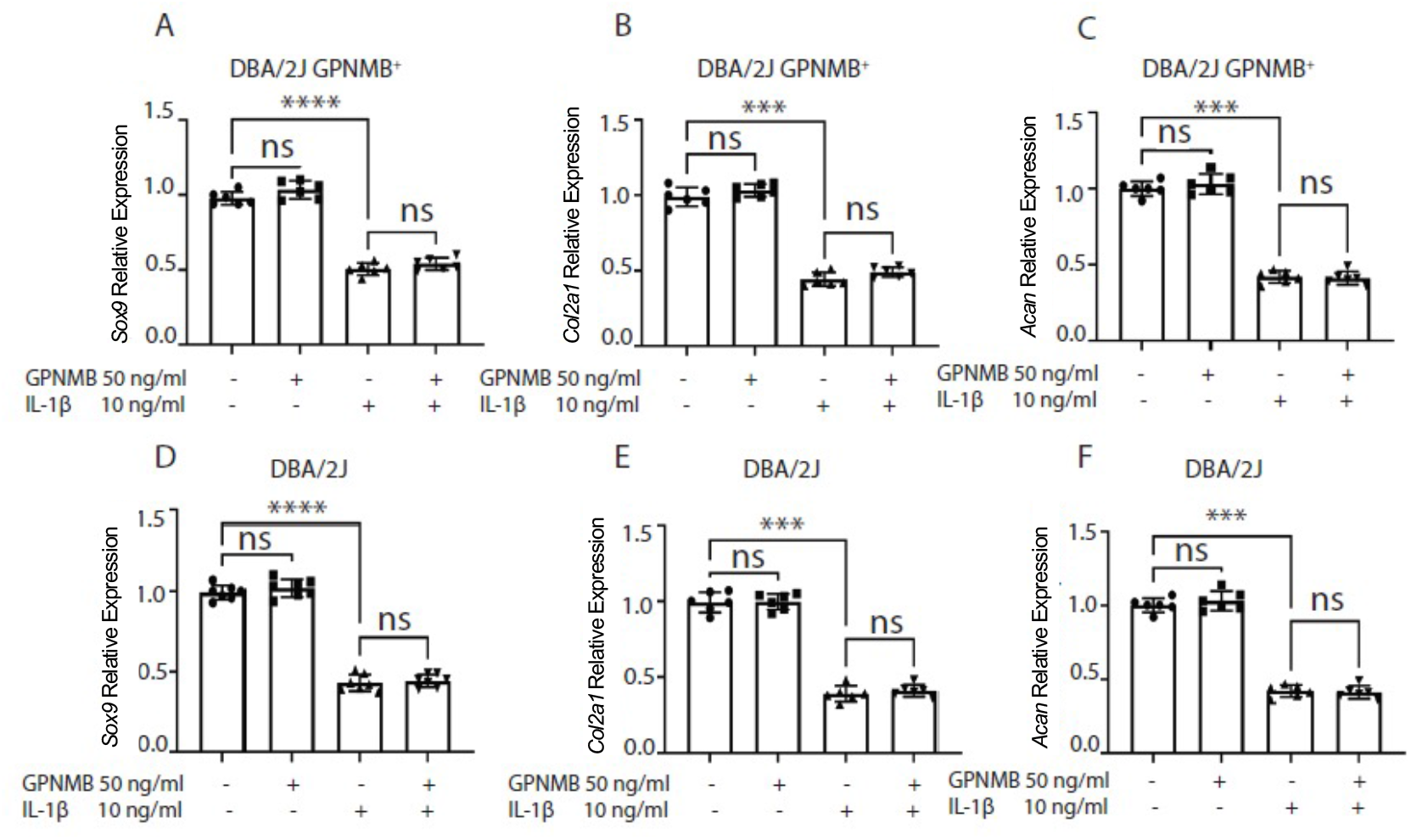
Recombinant GPNMB treatment does not alter anabolic gene expression. Anabolic gene expression was measured in DBA/2J GPNMB^+^ and DBA/2J chondrocytes by RT-qPCR. *Sox9* mRNA was decreased with IL-1β treatment in DBA/2J GPNMB^+^ (**A**) and DBA/2J (**D**) chondrocytes and was not rescued following treatment with recombinant GPNMB. *Col2a1* expression was decreased with IL-1β treatment and was not increased significantly upon addition of recombinant GPNMB in DBA/2J GPNMB^+^ (**B**) or DBA/2J (**E**) chondrocytes. *Acan* mRNA expression decreased with IL-1β treatment and was not rescued with the addition of recombinant GPNMB in DBA/2J GPNMB^+^ (**C**) or DBA/2J (**F**) chondrocytes. Data presented as Mean ± SEM (n=6, 3 replicates per experiment). ns= not significant, **= p<0.01, ***=p<0.001, ****=p<0.0001.

Data from the DBA/2J chondrocytes (n=6) also showed no significant differences between untreated control cells and those treated with recombinant GPNMB alone (**Figure 3D, E, F**), while treatment with IL-1β alone significantly reduced expression of *Sox9, Col2a1* and *Acan* (**Figure 3D, E, F**). The combination treatment of IL-1β and recombinant GPNMB also failed to rescue anabolic gene expression in DBA-2J chondrocytes (**Figure 3D, E, F**). These results confirm that IL-1β stimulation strongly inhibits anabolic gene expression and demonstrates that recombinant GPNMB treatment alone is not sufficient to rescue anabolic gene expression in DBA/2J GPNMB^+^ or DBA/2J primary chondrocytes.

### DBA/2J mice display an accelerated progression of articular cartilage degeneration

The *in vitro* experiments demonstrated recombinant GPNMB treatment inhibited IL-1β-induced inflammation. To understand the possible role GPNMB may play in chondrocytes *in vivo*, we utilized a post-traumatic OA surgical mouse model, destabilization of the medial meniscus (DMM) (Culley et al., 2015; Glasson et al., 2007). First, we confirmed successful DMM surgery in C57BL/6J mice (n=7). Mice were subjected to DMM joint surgery to induce OA on the right joint and sham surgery on the left, which served as internal controls. The mice were sacrificed at 12-weeks post-surgery and joints were histologically analyzed via Toluidine Blue and Safranin O staining (**Figure 4 A-B**). The severity of the damage was analyzed following the Osteoarthritis Research Society International (OARSI) (**Figure 4C**) (Moskowitz, 2000; Pritzker et al., 2006). After 12 weeks, cartilage was intact in sham operated joints and in DMM operated joints fibrillation and erosion was visible in superficial and intermediate layers indicating our DMM surgery was successful.

**Figure 4:**
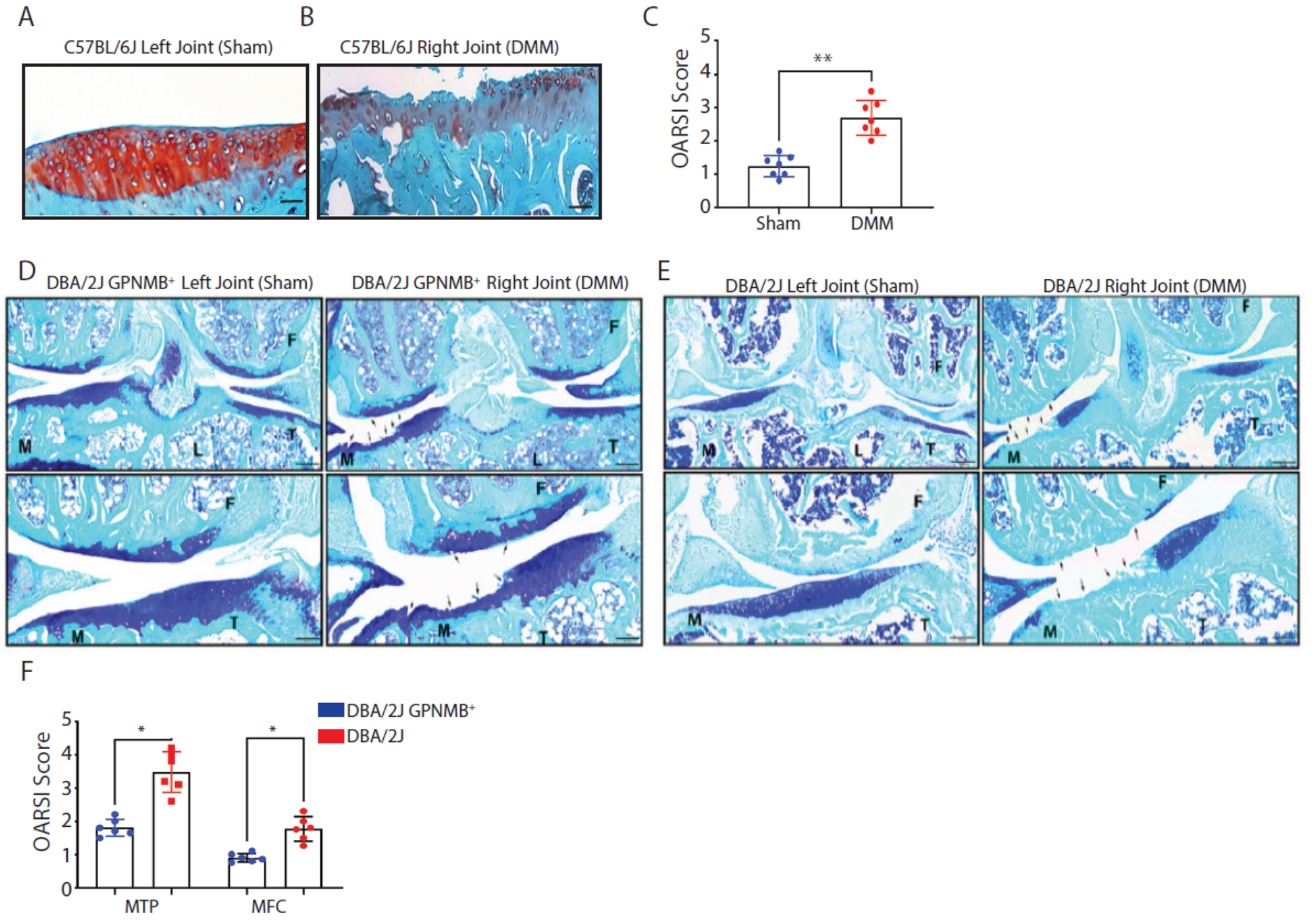
DMM surgery resulted in cartilage fibrillation after 12 weeks in C57BL6/J mice and damage was accelerated in DBA/2J mice following DMM. DMM surgery was validated in C57BL/6J mice. Cartilage was intact in sham operated joints (**A**) while DMM joints displayed superficial and intermediate cartilage damage (**B**) (tibia shown). (**C**) OARSI scores for sham and DMM operated joints. DBA/2J GPNMB^+^ (**D**) and DBA/2J (**E**) mice, which lack functional GPNMB protein, were subjected to sham and DMM surgery and joints were stained with Toluidine blue. Black arrows indicate damaged areas. (**F**) OARSI scores for femur (MFC) and tibia (MTP) cartilage damage of DBA/2J GPNMB^+^ and DBA/2J mice. Scale bar= 200µm in C57BL/6J joints and top panels and 100µm in magnified lower images (**D, E**). Images shown are representative and OARSI scorings from multiple images of n=7 each C57BL/6J, DBA/2J GPNMB^+^ and DBA/2J mice. Data presented as Mean ± SEM. *p<0.05, **= p<0.01.

Next, DBA/2J GPNMB^+^ and DBA/2J (n=7 each strain) mice were subjected to DMM surgery as described above. Cartilage damage caused by joint instability was observed in the articular cartilage of the medial compartment of the femur and tibia in both DBA/2J GPNMB^+^ and DBA/2J mice (**Figure 4D, E**). Compared to DBA/2J GPNMB^+^ control mice, the DBA/2J mutant mice demonstrated more severe cartilage damage (**Figure 4D, E**). The severity of the damage was analyzed following OARSI guidelines. DBA/2J GPNMB^+^ mice showed max scores of 1.629 ± 0.2616 for the medial tibial plateau (MTP) and 0.7571 ± 0.1043 for the femoral medial condyle (MFC) (**Figure 4F**). The DBA/2J mice were significantly higher at 2.656 ± 0.355 for the medial tibial plateau and 1.473 ± 0.202 for the femoral medial condyle (**Figure 4F**). These results show the GPNMB mutant mice, which lack a fully functional GPNMB protein, displayed more severe cartilage damage upon DMM surgery compared to the controls.

### DBA/2J exhibit reduced ACAN and enhanced IL-6 expression compared to DBA/2J GPNMB^+^

Given the severity of the cartilage damage seen in DBA/2J mice following DMM surgery, we next determined if ACAN expression was negatively affected in the mutant mice via immunohistochemistry. ACAN was chosen for analysis since it is the major cartilage proteoglycan responsible for bearing compressive loads during perambulation (Roughley, 2001; Roughley & Mort, 2014). Sham operated joints showed intact cartilage and intact ACAN contents with no significant differences detected between the groups (**Figure 5A, B**). While the ACAN expression in DBA/2J GPNMB^+^ DMM joints was also relatively high, DMM joint levels were much lower relative to sham operated controls and much higher than ACAN levels in the DBA/2J DMM mouse joints that showed severe cartilage damage (**Figure 5A, B**). We then quantified percent positive cells and found no significant differences in ACAN positive cells in sham operated joints between strains (**Figure 5C**), but a significant difference in ACAN production between DMM joints of DBA/2J GPNMB^+^ and DBA/2J mice (**Figure 5D**). These results revealed that DBA/2J mice exhibited severe cartilage damage and that the damage is due, in part, to degradation of critical ECM components such as ACAN.

**Figure 5:**
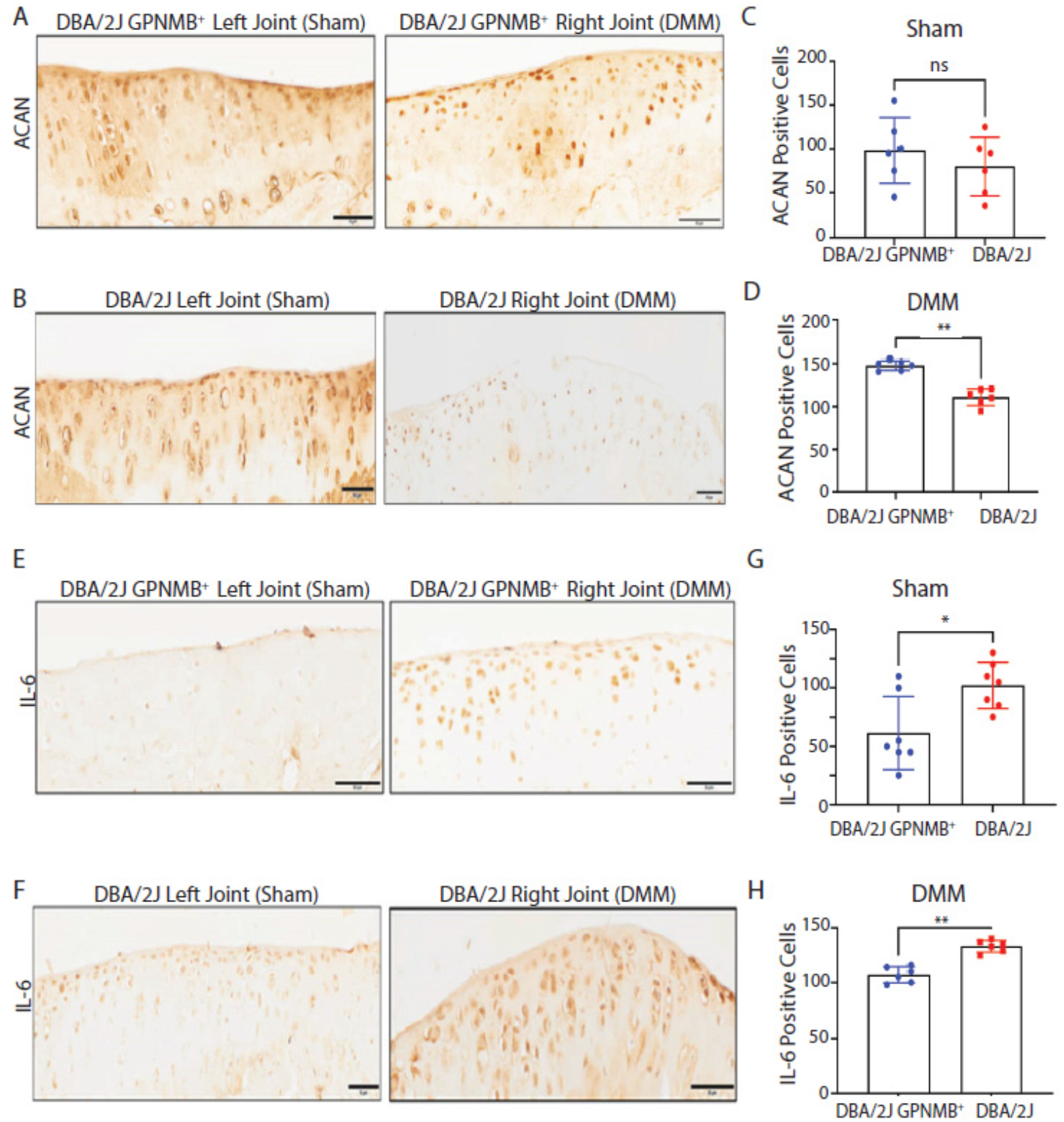
ACAN expression is severely depleted, and IL-6 expression elevated following DMM in DBA/2J mice. ACAN expression was not significantly altered in sham surgery mice (**A-C**) but was reduced following DMM in DBA/2J GPNMB^+^ (**A, D**) and was severely depleted in DBA/2J (**B, D**) cartilage. Endogenous IL-6 expression is increased in DBA/2J animals without DMM (**E-G**). DBA/2J GPNMB^+^ (**E, H**) and DBA/2J (**B, H**) cartilage showed an increase in IL-6 following DMM surgery. Scale bar= 200µm top panel and 100µm in magnified lower images. Images shown are representative and OARSI scores are taken multiple images of n=5-7 each DBA/2J GPNMB^+^ and DBA/2J mice. Data presented as Mean ± SEM. *p<0.05, **p<0.01.

In addition to the anabolic marker ACAN, we also evaluated the expression of the inflammatory marker IL-6 following DMM surgery in both DBA/2J GPNMB^+^ and DBA/2J mice (n=7 each strain). Sham and DMM joint sections were subjected to immunohistochemistry staining using an anti-IL-6 antibody. Both DBA/2J GPNMB^+^ and DBA/2J mice expressed IL-6 as an inflammatory response to the post-traumatic joint damage caused by DMM (**Figure 5E, F**). Quantification of IL-6 expression showed significantly higher number of IL-6 positive cells in DBA/2J mice following sham surgery compared to DBA/2J GPNMB^+^ controls (**Figure 5G**), suggesting DBA/2J mutant mice produce higher endogenous levels of proinflammatory IL-6 in the absence of functional GPNMB protein than control animals. This significance is further enhanced upon DMM surgery, where there is a significant increase in IL-6 positive cell numbers in the DBA/2J DMM joints compared to DBA/2J GPNMB^+^ control DMM results (**Figure 5H**).

### GPNMB inhibits MAPK Kinase signaling in mouse primary chondrocytes

Previous literature has shown that GPNMB might play a role in the mitogen-activated protein kinase (MAPK) pathway in a variety of cell types such as mesenchymal stem cells, fibroblasts, and osteoclasts (S. M. Abdelmagid et al., 2015; Furochi et al., 2007; Ono et al., 2016). In this study we determined the effects of recombinant Gpnmb on MAPK singling in primary chondrocytes. Primary chondrocytes were isolated from C57BL/6J male mice (n=9), treated with recombinant GPNMB with or without IL-1β and terminated at 30 and 50 minutes after treatment. Protein lysates were probed for extracellular signal-related kinase (ERK), phospho-ERK (pERK), and GAPDH for loading control. Data showed the pERK was increased following IL-1β stimulation and decreased upon recombinant GPNMB treatment (**Figure 6A, B**). Recombinant GPNMB treatment inhibited pERK after 30 minutes. These results were semi-quantified by densitometry analyses (**Figure 6C, D**). These results indicate recombinant GPNMB plays a role in pERK inhibition following inflammation induced by IL-1β stimulation.

**Figure 6:**
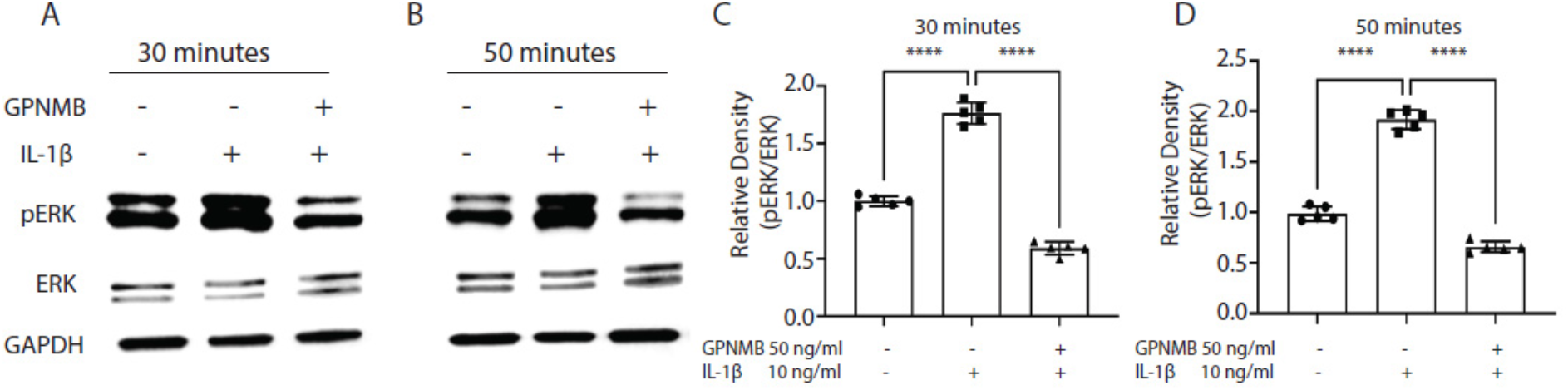
Recombinant GPNMB inhibits the MAPK signaling pathway in primary chondrocytes. Immunoblotting of primary chondrocytes following IL-1β stimulation with and without recombinant GPNMB show GPNMB inhibits pERK at (**A**) 30 and (**B**) 50 minutes post-treatment. Densitometric analyses of pERK were done at (**C**) 30 and (**D**) 50 minutes. Experiments were repeated three times with similar results. MW: pERK (44 kDa), ERK (42 kDa), GAPDH (37 kDa). Data presented as Mean ±SEM (n=5). ****=p<0.0001, ns= not significant.

## Discussion

While the role of GPNMB (osteoactivin) in fracture repair and bone regeneration is well characterized, nothing is known of its biological function in the joint or in chondrocytes, the principle cell type present in cartilage (Akkiraju & Nohe, 2015; Hall, 2019). Furthermore, while GPNMB expression is increased under inflammatory conditions in human and porcine chondrocytes, it is unclear what role GPNMB plays in the pathogenesis of cartilage degenerative disorders such as OA (Conde et al., 2015; Jiang et al., 2011; Owen et al., 2003; Schlichting et al., 2014; Sun et al., 2015). OA is characterized by articular cartilage degeneration, abnormal bone formation (osteophytes) and chronic inflammation of the joint that results in pain and loss of mobility in affected individuals (Goldring et al., 2008; Goldring & Otero, 2011; Goldring et al., 2011; Goldring & Goldring, 2016; Loeser et al., 2016; Loeser et al., 2012). GPNMB has been shown to have anti-inflammatory effects in lymphocytes, macrophages, periodontal ligament cells and in microglia and astrocytes (Budge et al., 2020; Li et al., 2010; Neal et al., 2018; Ripoll et al., 2007; Song & Lin, 2019; Zhou et al., 2017). Given these findings, we examined the biological role of GPNMB on inflammation and cartilage damage in primary chondrocytes *in vitro* and in articular cartilage *in vivo*. We provide the first evidence that GPNMB protects against cartilage damage in a post-traumatic mouse model and inhibits the expression of inflammatory cytokines and chemokines without altering anabolic gene expression in human and mouse primary chondrocytes.

In humans, *GPNMB* was increased in damaged high-grade osteoarthritic cartilage, primary chondrocytes, and chondrocyte cell line *in vitro* and expression was increased with IL-1β proinflammatory stimulation. Additionally, treatment with recombinant GPNMB significantly reduced catabolic gene expression in primary human and mouse chondrocytes without altering anabolic gene expression, suggesting GPNMB does not affect chondrogenesis. Additionally, chondrocytes derived from mice with reduced levels of GPNMB had significantly higher expression of catabolic and pro-inflammatory genes following inflammatory stimulation compared to control chondrocytes. This suggests that endogenous Gpnmb may play a role in modulating inflammation.

Expanding on our *in vitro* findings, we next determined the role of GPNMB *in vivo* via a DMM model of induced post-traumatic OA in mice (Culley et al., 2015; Glasson et al., 2007). Surgeries were performed on DBA/2J GPNMB^+^ controls and in DBA/2J mice, which produce a non-functional truncated version of the GPNMB protein (Abdelmagid et al., 2014; Chung et al., 2007; Gutknecht et al., 2015). Inactive GPNMB leads to increased immune function and inflammation in these animals, with proinflammatory cytokine levels consistently higher than DBA/2J GPNMB^+^ controls (Fan et al., 2010; Wilson et al., 2015; Zhou et al., 2009). When cartilage integrity was assessed post-surgery, DBA/2J mice displayed severe cartilage damage, suggesting GPNMB is beneficial for cartilage homeostasis.

To determine the signaling pathway responsible for the GPNMB-mediated reduction of inflammation in cartilage and primary chondrocytes, we evaluated the effects of GPNMB on the MAPK signaling pathway which is required for chondrocyte expression of MMPs, inflammatory cytokines, and ADAMTS in cartilage (Cui et al., 2020; Goldring, 1999; Tetsunaga et al., 2011). GPNMB is known to function via the ERK pathway in osteoclasts, motor-neuron-like cells, and fibroblasts (Furochi et al., 2007; Ono et al., 2016; Sondag et al., 2016). Also, ERK pathway activation has been shown to regulate osteoblast proliferation and differentiation (Bellido & Plotkin, 2011; Jessop et al., 2002; Yan et al., 2012). However, in chondrocytes, activation of the ERK pathway has been shown to be detrimental resulting in chondrodysplasia and contributing to cartilage degradation in the pathogenesis of OA (Appleton, 2018; Appleton et al., 2010; Lin et al., 2021; Yang et al., 2008). Here, we are the first to show that the anti-inflammatory action of GPNMB in chondrocytes occurs via MAPK in primary chondrocytes with IL-1β-induced inflammation (**Figure 7**). Furthermore, our results suggest the strong anti-inflammatory function of GPNMB may be a result of concurrent inhibition of multiple catabolic markers (MMP-3, MMP-9, MMP-13, ADAMTS-4, and IL-6).

**Figure 7:**
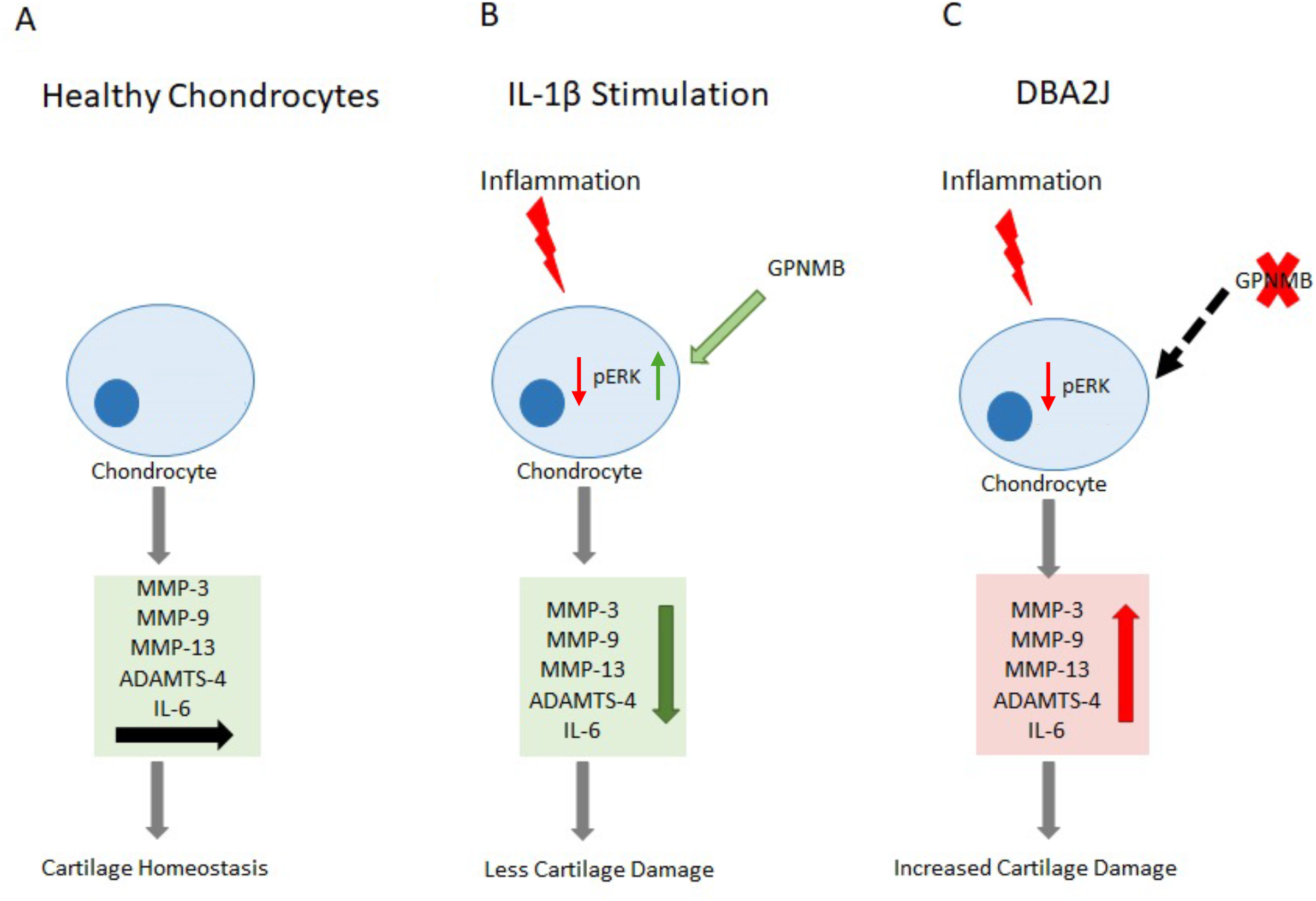
Schematic of the proposed mechanism of GPNMB anti-inflammatory function in chondrocytes. (**A**) In healthy chondrocytes, GPNMB functions without inflammatory stimuli to modulate catabolic chemokines and cytokines and maintain cartilage homeostasis. (**B**) In the presence of inflammatory stimuli (IL-1β), GPNMB acts to modulate ERK to reduce expression of catabolic chemokines and cytokines and reduce cartilage damage. (**C**) In DBA/2J mice with non-functional GPNMB, there is no inhibition of inflammatory molecules leading to increased catabolic expression and increased cartilage damage.

In conclusion, the data presented here demonstrates a novel anti-inflammatory role for GPNMB in cartilage and primary chondrocytes. While more work is needed to fully elucidate the mechanism *in vitro* and *in vivo*, these novel findings show GPNMB reduces catabolic gene expression and lays the foundation for future studies to assess the use of GPNMB as a possible alternative therapeutic agent in the treatment of OA.

## Materials and Methods

### HTB-94 Cell Line Culture and Treatment Conditions

HTB-94 cells were purchased and authenticated by American Type Culture Collection (SW1353). Cells were then cultured in DMEM/F12 (1:1, Cellgro Technologies, Lincoln, NE) culture medium supplemented with Penicillin/Streptomycin (Penicillin: 100 IU/ml, Sigma Aldrich, St. Louis, MO; Streptomycin: 100 µg/ml, Sigma Aldrich) and 10% fetal bovine serum (FBS, GE Healthcare, Chicago, IL) at 37°C and 5% CO_2_. Once established in culture, monolayer HTB-94 cells were rinsed with sterile PBS and serum-starved overnight prior to stimulation with 10 ng/ml of recombinant interleukin-1β (IL-1β; R&D Systems, Niskayuna, NY) with or without recombinant GPNMB (R&D Systems) in DMEM/F 12 (1:1) medium supplemented with Penicillin (100 IU/ml, Sigma-Aldrich), Streptomycin (100 μg/m, Sigma-Aldrich). HTB-94 experiments were set up in triplicate and results shown are from three separate experiments.

### Human Explant Culture and GPNMB/IL-1β Treatment

Explants were obtained from undamaged regions of discarded and de-identified articular knee cartilage harvested from patients following knee arthroplasty (n=7). Undamaged cartilage regions were again identified using India Ink and then washed twice in sterile PBS. Cartilage explants (2 x 2 mm) were excised using a biopsy punch and maintained for 24 hours in supplemented DMEM/F12 culture medium as described above at 37°C and 5% CO_2_. After 24 hours, the medium was replaced with serum free DMEM overnight and then treated with 10 ng/ml IL-1β with or without recombinant 50 ng/ml GPNMB for 24 hours. Following pharmacological treatment, the explants were fixed in 4% paraformaldehyde (Sigma Aldrich) for 24 hours, decalcified using 14% Ethylenediaminetetraacetic acid (EDTA, Thermo Fisher Scientific, Waltham, MA), and processed. Explants were sectioned (5µm) and stained with Safranin-O stain (Thermo Fisher Scientific) (Schmitz et al., 2010).

### Mouse Strains

The following mouse strains were utilized in the completion of this study: C57BL/6J (wild type, WT, Stain #: 000664), DBA/2J (GPNMB mutant, Strain #: 000671), and DBA/2J GPNMB^+^ (GPNMB control, Strain #: 007048). Mice were purchased from Jackson Laboratories and were housed at NEOMED in accordance with protocol guidelines approved by the NEOMED Institutional Animal Care and Use Committee (IACUC) under protocol 23-10-385. The DBA/2J GPNMB+ mice are a coisogenic strain purchased from Jackson Laboratories. Jackon Laboratories generated these mice by knocking in the wild-type allele of Gpnmb into the DBA/2J background. By doing so, they rescued the phenotype of the DBA/2J mice. This description has been highlighted in our previous publications (Abdelmagid et al., 2014; Samir M. Abdelmagid et al., 2015).

### Murine Primary Chondrocyte Isolation and Culture and Treatment Conditions

Primary articular chondrocytes isolated from hind limb dissections of 5-6-day-old mouse pups as previously described (Gosset et al., 2008; Jonason et al., 2015). In brief, skin and soft tissues were removed from the hindlimb bones and the femoral head, femoral condyle and tibial plateau cartilage were removed and placed in 1X sterile PBS on ice. Cartilage was washed twice with cold 1X sterile PBS and incubated twice (45 minutes per incubation) in a digestion solution consisting of Collagenase D (3 mg/ml, Sigma Aldrich) in serum free Dulbecco’s modified Eagle’s medium (DMEM) supplemented with Penicillin (100 IU/ml) and Streptomycin (µg/ml) at 37°C and 5% CO_2_. Then, cartilage digestion solution was diluted with serum free culture medium to a final concentration of 0.5 mg/ml and the cells were incubated overnight. After digestion, the cell solution was re-suspended and filtered using a sterile 40 µm cell strainer (Thermo Fisher Scientific). The filtered cell solution was centrifuged at 400g for 10 minutes, rinsed twice with sterile 1X PBS (centrifuging after each rinse), and re-suspended in culture medium consisting of DMEM supplemented with Penicillin (100 IU/ml), Streptomycin (100 μg/m) and 10% FBS. Mouse primary chondrocytes were counted via hemocytometer and seeded at high density in a 6-well culture dish for four days. Chondrocytes were serum stared overnight on day 5 and treated on day 6 with recombinant IL-1β (10 ng/ml) and recombinant GPNMB (50 ng/ml). On day 7, cells were harvested for immunoblotting and total RNA extraction.

### Cell Viability and Proliferation Assays

Cell viability in response to different doses of recombinant GPNMB was assessed using the MTT assay (Thermo Fisher Scientific). One day prior to treatment, primary chondrocytes were seeded into 96-well culture plates. Chondrocytes were treated for 72 hours with varying doses of recombinant GPNMB (R&D systems) and the culture media was aspirated and 20 µl of MTT solution was added to the cells for 4 hours at 37°C and 5% CO_2_. Then, the supernatant was aspirated, dimethyl sulphoxide was added with gentle agitation for 15 minutes and absorbance was measured at 570 nm on an ELISA BioTek plate reader (BioTek, Winooski, VT). Cell viability assays were conducted 3 times (3 replicates per group) and treatment results were compared to untreated control cells.

Cellular proliferation of primary chondrocytes was determined with the CyQuant NF cell proliferation assay (Life Technologies, Carlsbad, CA). Mouse primary chondrocytes were harvested from 45-day-old pups and isolated as previously described. Cells were seeded in a 96-well culture plate for 24 hours and then treated with different concentrations of recombinant GPNMB for 72 hours. The media was then aspirated and replaced with 100 µl 1X binding solution and incubated for 1 hour at 37°C. Fluorescent intensity was measured on a BioTek plate reader (excitation at 485 nm and emission at 530 nm). Proliferation assays reported are the results of 3 separate experiments (3 replicates per group) and results of treated cells were compared to those of untreated control cells.

### RNA Isolation and RT-qPCR Analysis of Gene Expression

Total RNA was isolated from primary chondrocytes using Qiazol (Qiagen, Hilden, Germany) and purified using the RNeasy Mini Kit (Qiagen) following the manufacturer recommended protocols. RNA was quantified on a Nanodrop 2000 spectrophotometer (Nanodrop, Thermo Fisher Scientific). cDNA was synthesized from 1 µg total RNA using the High-Capacity cDNA Reverse Transcription kit (Thermo Fisher Scientific).

RT-qPCR analyses were completed in triplicate using in-house designed primers (**Table 1**) and SYBR green PCR Master Mix (Thermo Fisher Scientific) on a Step One Plus Machine (Applied Biosystems, Waltham, MA). Expression of target genes was determined utilizing the delta-delta-Ct method and GAPDH was employed for normalization as an endogenous control (Livak & Schmittgen, 2001; Schmittgen et al., 2008; Schmittgen & Livak, 2008).

**Table 1:**
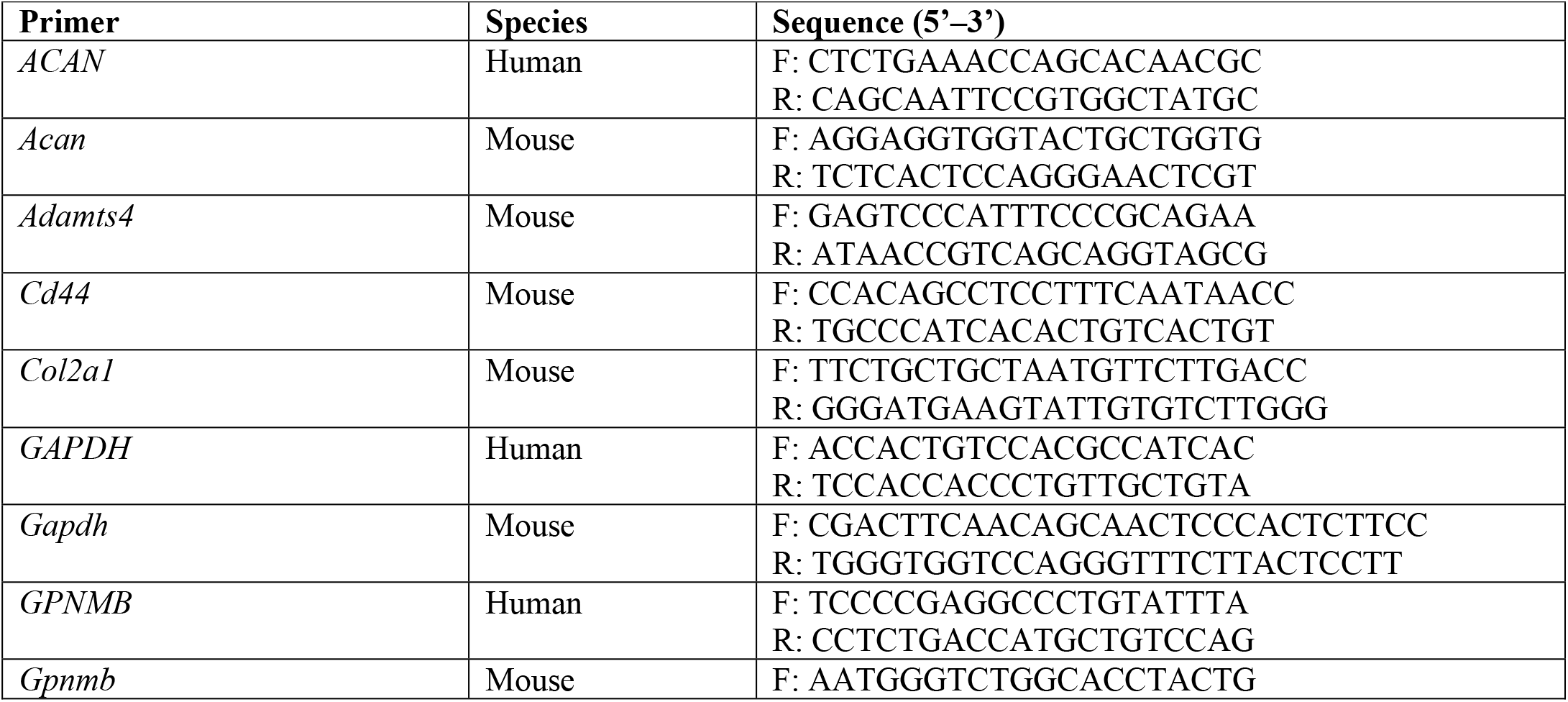
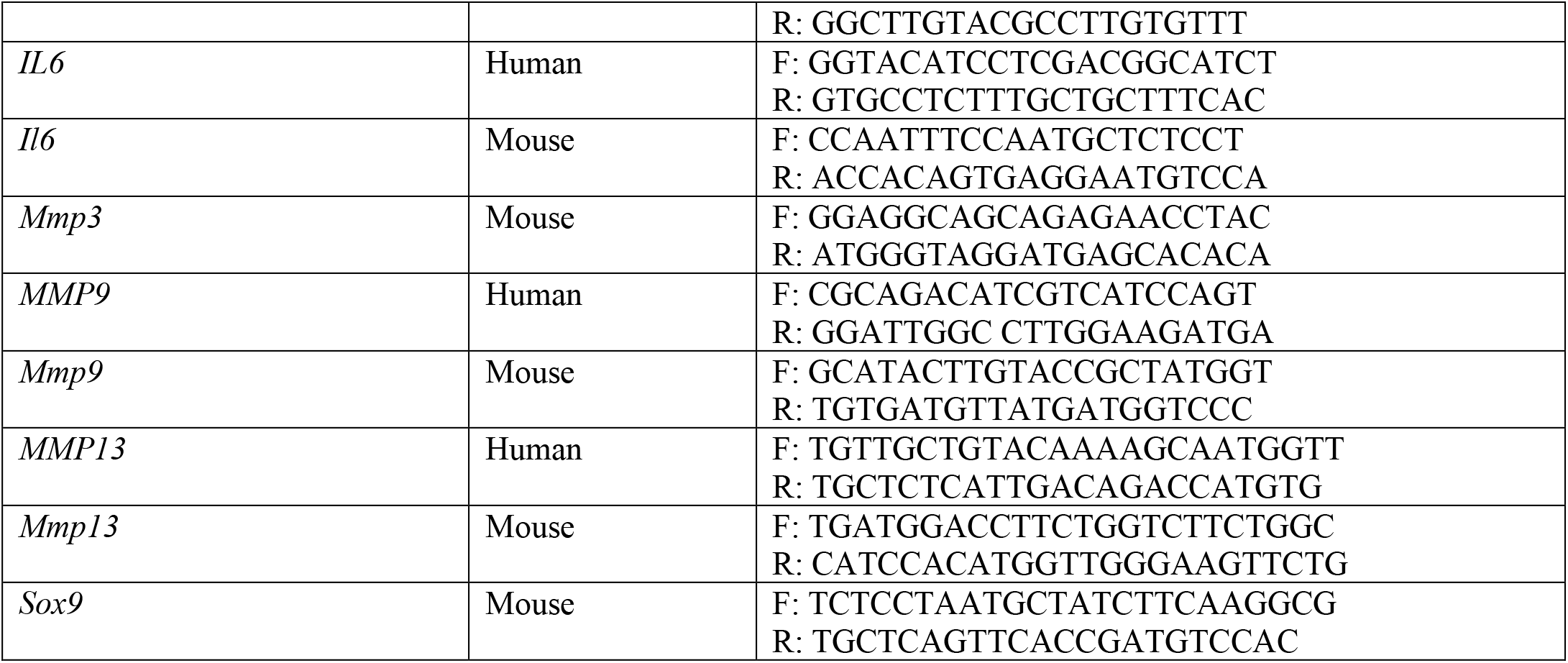
qPCR primers.

### Immunoblotting

Total cellular protein was isolated from primary chondrocytes using radioimmunoprecipitation buffer (RIPA, Millipore, Burlington, MA) with added phosphatase inhibitor (Pierce, Rockford, IL). Protein concentrations were established using the Pierce BCA Protein Assay Kit (Pierce). Equivalent protein concentrations (20 µg) were loaded with 2X sample buffer (BioRad, Hercules, CA) into 12% SDS-PAGE gels. The Trans-Blot Turbo System (BioRad) was used to transfer proteins to a polyvinylidene fluoride membrane (PVDF, BioRad). Membranes were then blocked with 5% bovine serum albumin (BSA, Cytiva, Marlborough, MA) in 0.1% Tri-Buffered Saline with Tween 20 (TBST) for 1 hour at room temperature and then incubated as described at 4°C overnight with primary antibodies listed in **Table 2**. Membranes were rinsed twice in TBST (5 minutes per wash) with gentle shaking and a Horseradish peroxidase (HRP)-linked secondary in TBST (Cell Signaling Technologies, Danver, MA) was applied to the membranes for 1 hour. Immunoblots were developed with chemiluminescent substrate (Pierce). Bands images and intensity were determined using a Pxi imaging system and Syngene software (Syngene, Bangalore, India).

**Table 2:** Table of Primary Antibodies.

| Source | Primary Antibodies | Company | Catalog Number | RRID |
| --- | --- | --- | --- | --- |
| Rabbit | ACAN | Bioss | BS-11655R | N/A |
| Rabbit | ERK | Cell Signaling Technologies | 9102 | AB_330744 |
| Rabbit | pERK | Cell Signaling Technologies | 9101 | AB_331646 |
| Rabbit | GAPDH | Cell Signaling Technologies | 3683 | AB_1642205 |
| Rabbit | IL-6 | Abcam | AB6672 | AB_2127460 |

### DMM Surgery

DMM was performed in 12-week-old male DBA/2J GPNMB^+^ (control, n=7) and DBA/2J (mutant, n=7) mice to induce OA. DMM was performed as previously described and surgical procedures were approved by the NEOMED IACUC committee (Culley et al., 2015; Glasson et al., 2007). Only male mice were studied at this time. Although recent literature has shown that female mice do in fact develop post traumatic-OA, due to the preliminary nature of this study and resource constrains, only male mice were studied at this time (Hwang et al., 2021; Ma et al., 2007). Female mice will be included in future studies. The right knee joint was subjected to microsurgery under a sterilized microscope. The medial menisci tibial ligament (MMTL), which anchors the medial meniscus (MM) to the tibial plateau, was transected to destabilize the joint. The left knee joint was subjected to sham surgery as a control. The knee was surgically opened but the MMTL was not transected. Mice were sacrificed 12-weeks post-surgery.

### DMM Tissue Isolation

The sham and DMM surgery mouse knee joints were harvested as previously described (Culley et al., 2015). Soft tissues were removed and the joints were fixed for 24 hours in 4% paraformaldehyde with gentle agitation. The fixed joints were rinsed for 1 hour in PBS and then decalcified for 10 days in 14% EDTA. The buffer was changed three times per week during the decalcification process. Decalcified joints were washed for 6 hours in PBS, processed, and embedded in paraffin and cut into 6 µm sections.

### Histological Assessment of Mouse Sham and DMM Knee Joints

To assess cartilage damage in the sham and DMM surgery mouse joints, the sections were stained with Toluidine blue/fast green stain. Three sections were selected at 90 µm intervals for histological evaluation following the Osteoarthritis Research Society International (OARSI) scoring system (Pritzker et al., 2006). Results were presented as ± mean of the maximum joint score.

### Immunohistochemistry

Histological sections of sham and DMM mouse knee joints were subjected to immunohistochemical analysis to measure expression of catabolic and anabolic proteins in damaged and undamaged regions of the cartilage. Histological sections were de-paraffinized for 1 hour and cleared in xylene (Thermo Fisher Scientific) before they were rehydrated in a graded ethanol series. For antigen retrieval, the sections were treated with 0.01 M citrate buffer at 60°C overnight in a water bath. Activity of endogenous peroxidase was quenched using 3% hydrogen peroxide (H_2_O_2_) solution for 15 minutes before blocking with 5% BSA for 1 hour. Sections were then treated overnight at 4°C with primary rabbit anti-Mmp-13 (Abcam, Cambridge, England), rabbit anti-Il-6 (Abcam), rabbit anti-Gpnmb (LSBio, Lynnwood, WA) or rabbit anti-Acan (Bioss, Woburn, MA). Biotinylated goat anti-rabbit secondary antibody (Vector Laboratories, Newark, CA) was added for 1 hour and the Vectastain Elite ABC HRP kit was sued with 3, 3’-Diaminobenzidine (DAB, Vector Laboratories) was used for signal development. The sections were imaged using a BX61VS microscope (Olympus Life Science, Center Valley, PA) and positive cells were quantified with ImageJ software (Schneider et al., 2012).

### Statistical Analyses

For all data generated, differences between individual groups were analyzed using GraphPad Prism software (GraphPad, La Jolla, CA). All cell culture experiments were repeated a minimum of three times with similar results. DMM experiments were repeated twice with 6 individuals included in each trial. In comparisons of multiple groups, One-way analysis of variance (ANOVA) tests were employed followed with Tukey’s post hoc tests. Comparisons of two groups employed unpaired *T*-tests. For histological scoring, MannWhitney and KruskalWallis tests were performed with Dunn’s post hoc analyses were conducted as the nonparametric equivalents for the independent samples t-test and One-way ANOVA. Differences were considered significant where p<0.05 and graphs depict group means + standard error of the average (±SEM).

## Acknowledgements

Research reported in this publication was supported by a generous gift from the Cook Family Fund for Orthopaedic Research, the National Institute Of Arthritis And Musculoskeletal And Skin Diseases of the National Institutes of Health under Award Number F30AR085476, the National Institute of Aging of the National Institutes of Health, Small Business Innovation Research Grant, under Award Number R43AG071396, and Biomedical Sciences Graduate Program at Kent State University. The content is solely the responsibility of the authors and does not necessarily represent the official views of the National Institutes of Health.

## Data and Materials Availability

No newly created dataset were generated in this manuscript.

---

**Figure 1-Figure Supplement 1:**
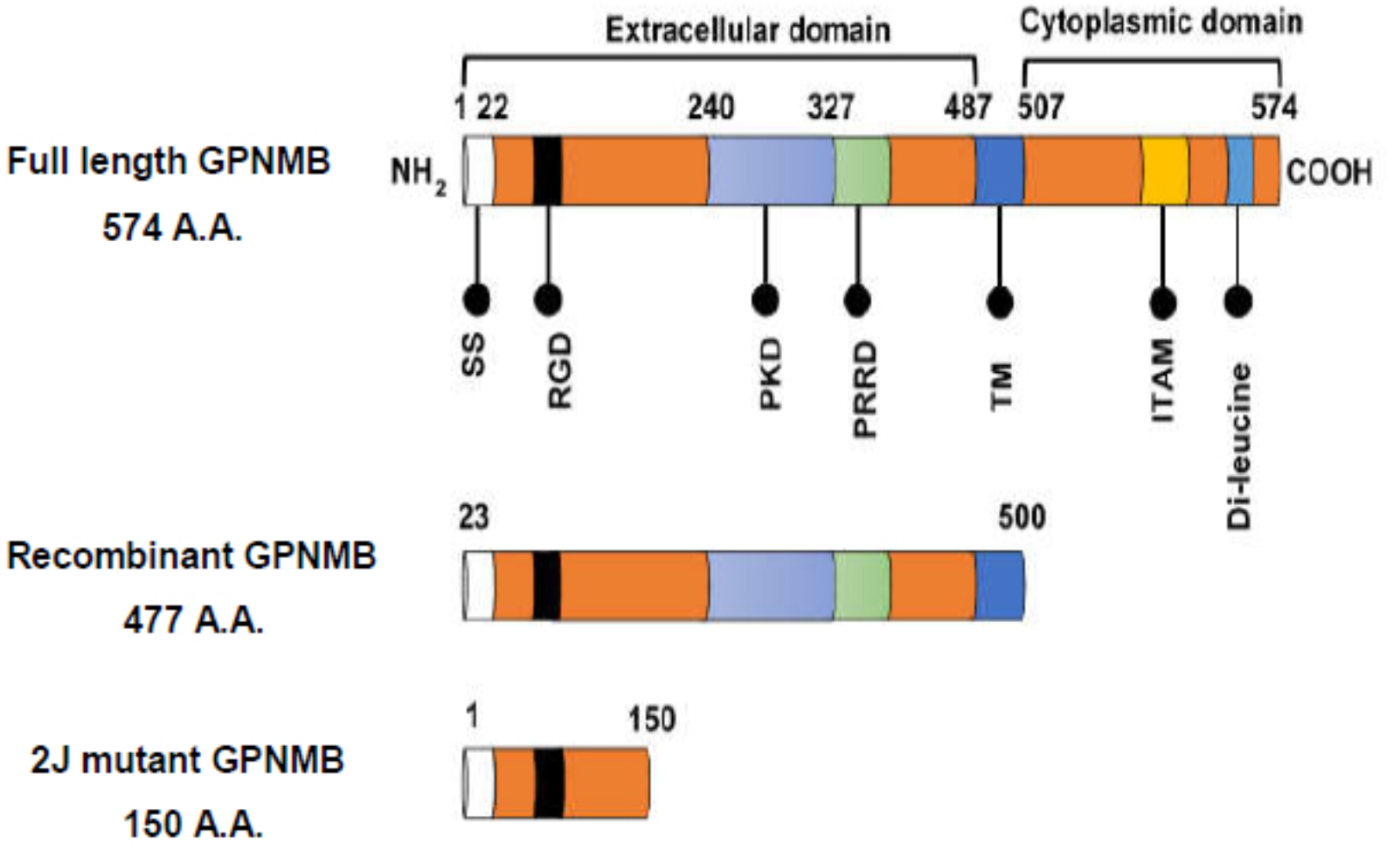
The structure of full length, recombinant and mutant GPNMB proteins. The full length GPNMB is 574 amino acids and has several domains including a N-terminal extracellular domain signal peptide (SS, 1-22), an arginine-glycine-aspartic acid (RGD), a polycystic kidney disease (PKD), and a proline rich repeat (PRRD) domain. The transmembrane domain (TM) of GPNMB is 20 amino acids (503-523). The C-terminal intracellular domain holds an ITAM motif and Di-leucine signal. The recombinant form of GPNMB represents the N-terminal portion of the molecule (23-502) and is derived from a mouse myeloma cell line. The DBA/2J mutant GPNMB is a truncated version of the protein (1-150).

**Figure 2-Figure Supplement 2:**
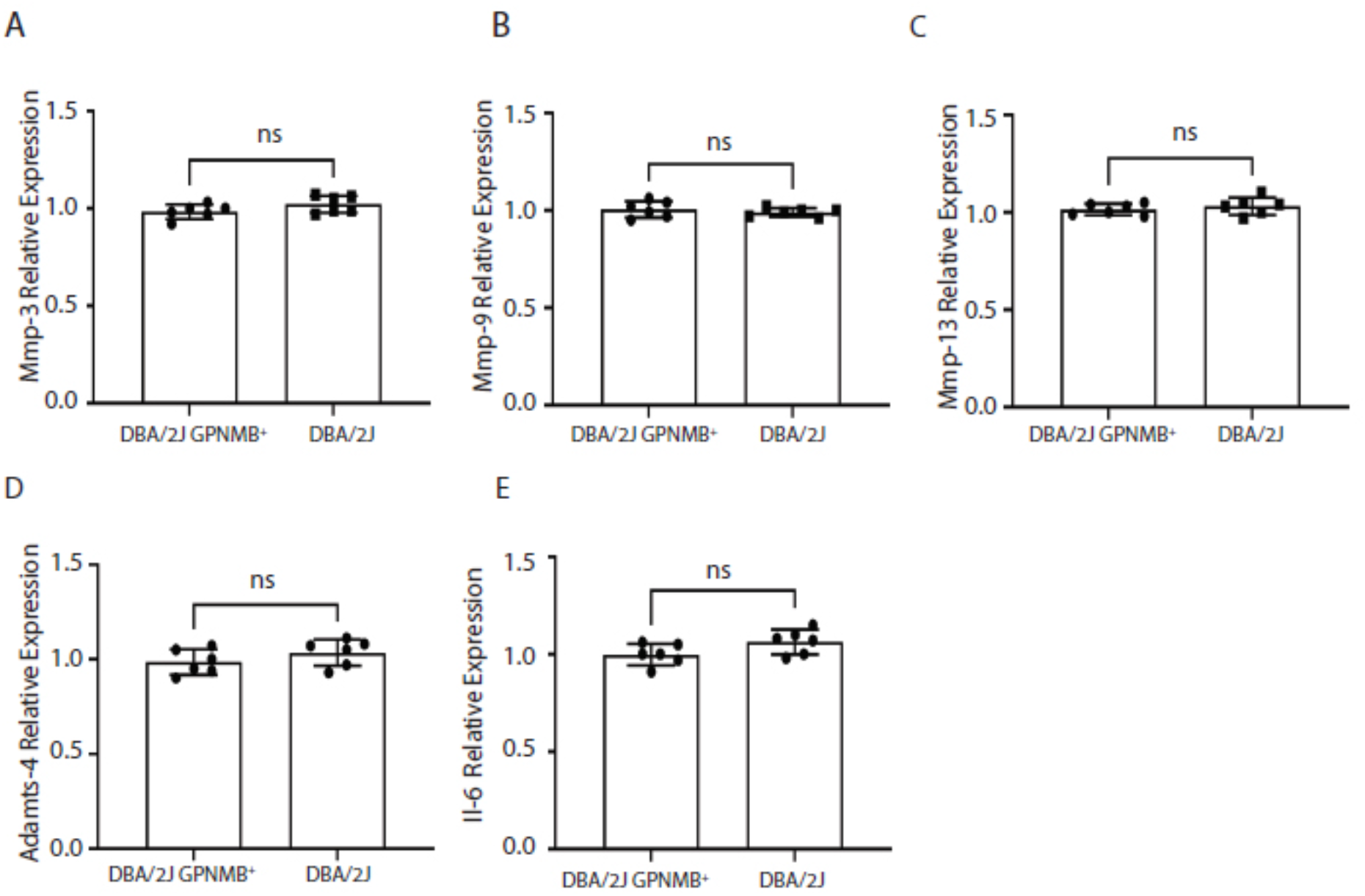
Endogenous levels of catabolic genes are not significantly different between DBA/2J GPNMB^+^ and DBA/2J primary chondrocytes. RT-qPCR analyses of (**A**) Mmp-3, (**B**) Mmp-9, (**C**) Mmp-13, (**D**) Adamts-4 and (**E**) Il-6 expression detected no significant differences in endogenous mRNA expression prior to treatment. Data presented as Mean ± SEM (n=6, 3 replicates per experiment); ns = not significant.

**Figure 3-Figure Supplement 3:**
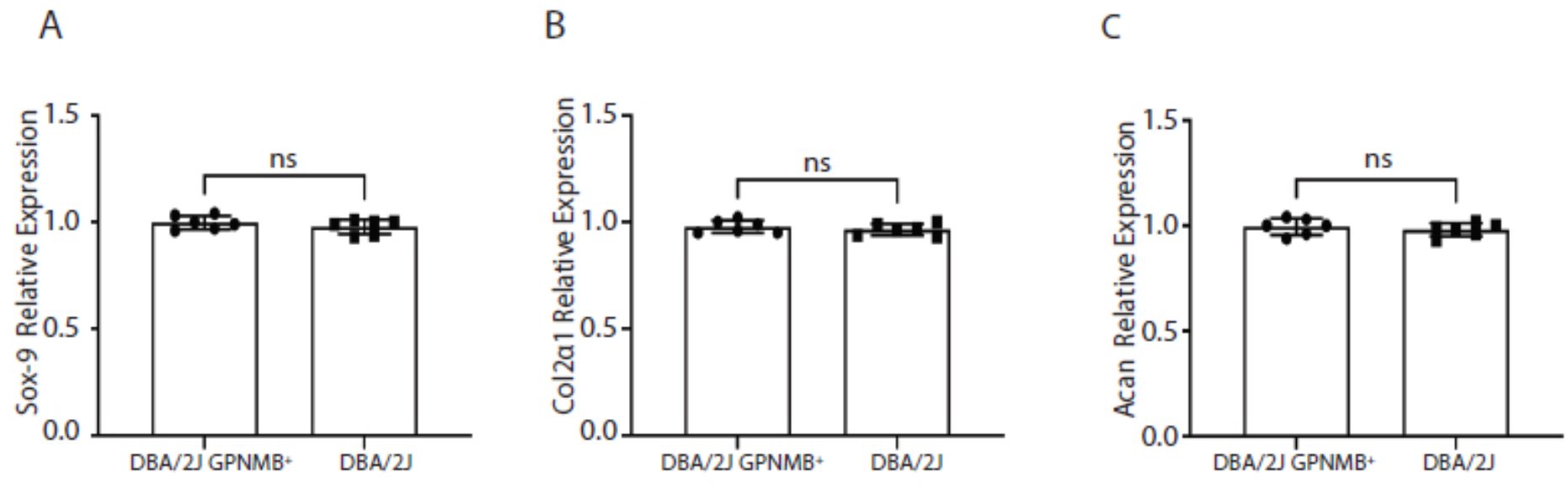
Endogenous levels of anabolic genes are not significantly different between DBA/2J GPNMB^+^ (control) and DBA/2J primary chondrocytes. RT-qPCR analyses of endogenous anabolic gene expression found no significant differences in (**A**) Sox-9, (**B**) Type-II collagen (Col2α1) or (C) ACAN expression. Data presented as Mean ± SEM (n=6, 3 replicates per experiment); ns = not significant.

